# Comprehensive interactome analysis for the sole adenylyl cyclase Cyr1 of *Candida albicans*

**DOI:** 10.1101/2022.03.15.481403

**Authors:** Guisheng Zeng, Suat Peng Neo, Li Mei Pang, Jiaxin Gao, Jayantha Gunaratne, Yue Wang

## Abstract

Cyr1, the sole adenylyl cyclase of the fungal pathogen *Candida albicans*, is a central component of the cAMP/PKA signaling pathway that controls the yeast-to-hyphal morphogenesis. Cyr1 functions as a multivalent senor and integrator of various external and internal signals. To better understand how these signals are relayed to Cyr1 and how Cyr1 activity is regulated, we sought to identify global interacting partners of Cyr1 using stable isotope labeling by amino acids in cell culture (SILAC)-based quantitative proteomics. From the proteins co-immunoprecipitated with Myc-tagged Cyr1, 38 were assigned as authentic Cyr1-interacting partners, including two previously characterized Cyr1-binding proteins, Cap1 and Act1. Many of the identified proteins remain uncharacterized, but some can be classified into several functional groups, such as actin regulatory proteins, cell wall components, and mitochondrial proteins. We used biochemical and genetic methods to further characterize the interaction of Cyr1 with Mp65, a cell surface mannoprotein previously reported to have a role in cell wall integrity and possess several virulence-related traits. Taken together, our comprehensive analysis of the Cyr1-interacting proteins provides important information for establishing the Cyr1 interactome and uncovers a potential role for cell wall proteins in Cyr1-mediated cAMP signaling.

## 1. Introduction

*Candida albicans* is a major opportunistic fungal pathogen of humans. It inhabits as a relatively harmless commensal in the microflora of the oral cavity and the gastrointestinal and urogenital tracts in most healthy people. However, this fungus can cause various mucosal infections in otherwise healthy individuals when conditions are favorable. In immunocompromised patients such as those undergoing cancer chemotherapy or tissue transplantation or suffering HIV infection, *C. albicans* can disseminate throughout the body to initiate life-threatening systemic diseases with high mortality [1,2].

*C. albicans* is a dimorphic fungus and can proliferate in either a unicellular yeast or multicellular filamentous form. It grows as budding yeast-like cells under conditions of moderate temperature and low pH. In response to a range of environmental stimuli such as temperature elevation to 37°C, neutral pH, and exposure to serum, N-acetylglucosamine, and peptidoglycan, the yeast cells quickly switch to the filamentous growth mode forming pseudohyphae or true hyphae [3–5]. The ability of *C. albicans* to transition between yeast and filamentous forms of growth is closely correlated with its pathogenicity, as mutants locked in either form of growth were found to be avirulent in a mouse model of systemic infection [6].

The yeast-to-filament growth switch of *C. albicans* is tightly controlled by multiple regulatory circuits, in particular the mitogen-activated protein kinase (MAPK) cascade and the cyclic adenosine monophosphate (cAMP)/protein kinase A (PKA) signaling pathway [7,8]. The cAMP/PKA pathway plays a dominant role in the serum-induced hyphal growth, as blocking this pathway, but not the MAPK pathway, abolishes true hyphal growth in response to serum induction [6,9–13]. A central component of the cAMP/PKA pathway is Cyr1 (also known as Cdc35), the sole adenylyl cyclase of *C. albicans* that catalyzes cAMP synthesis [9]. cAMP activates PKA, which activates the downstream transcriptional factor Efg1, leading to the expression of hypha-specific genes [7]. Cyr1 was first identified by its ability to functionally complement the conditional growth defect of *Saccharomyces cerevisiae* cells with mutations in the GTPases Ras1 and Ras2 [9]. *C. albicans cry1*Δ/Δ cells have no detectable intracellular cAMP and fail to undergo hyphal growth under all inducing conditions, but hyphal growth can be restored by adding exogenous cAMP to the medium [9]. Cyr1 is a large protein of 1690 amino acids containing multiple functional domains and has been proposed to function as a signal sensor and integrator in response to a diverse range of external and internal stimuli [14]. A putative Gα domain near the N-terminus of Cyr1 is thought to bind the G-protein α subunit Gpa2 to mediate glucose-induced cAMP signaling, although this interaction has not been demonstrated experimentally [15,16]. Next to the Gα domain is a Ras-association (RA) domain that interacts with the small GTPase Ras1 to regulate cellular cAMP levels [17]. The central region of Cyr1 contains a stretch of 14 leucine-rich repeats (LRR) capable of recognizing bacterial peptidoglycan (PGN) to stimulate cAMP production and promote hyphal growth [18]. The LRR domain may also mediate temperature sensing through interaction with the heat shock protein complex Hsp90/Sgt1 [19,20]. The C-terminal half of Cyr1 harbors the catalytic domain (CYC) that catalyzes cAMP synthesis from ATP. The CYC domain also carries a CO_2_ sensor to mediate CO_2_-induced filamentous growth [21]. A putative protein phosphatase 2C domain locates between the LRR and CYC domains, but its function remains unidentified [9]. At the extreme C-terminus of Cyr1 is a Cap1-binding domain that mediates the formation of a tripartite protein complex containing Cap1, Cyr1 and G-actin [22]. This complex has been proposed to sense the dynamic state of the actin cytoskeleton and thus influence the cyclase activity [22]. Despite the significant progress in dissecting the role of Cyr1, how this large multivalent protein functions as both signal sensor and integrator remains poorly defined. Therefore, we hypothesize that establishing and functionally characterizing the Cry1 interactome will provide crucial mechanistic insights.

The stable isotope labeling by amino acids in cell culture (SILAC)-based quantitative proteomics is a powerful tool to establish the interactome of a target protein [23]. SILAC is a strategy to label the whole proteome with amino acids metabolically, often lysine and arginine carrying stable heavy isotopes such as ^13^C or ^15^N, through protein synthesis and turnover during cell growth. When the same proteins from unlabeled (light) and labeled (heavy) samples are combined and analyzed together by high-resolution mass spectrometry (MS), pairs of chemically identical peptides of different stable-isotope compositions can be differentiated owing to their mass difference. The SILAC-based quantitative MS offers a comprehensive yet specific method to identify the interacting partners of a target protein. For example, cells expressing an epitope-tagged bait protein are grown in a medium with heavy amino acids, and cells expressing the epitope alone are grown in a medium with standard amino acids. After protein extraction and immunoprecipitation with an antibody against the epitope, the immunoprecipitates from the two cultures are combined for quantitative MS analysis. Nonspecific binding proteins will have an equal abundance from both light and heavy samples, while specific binding proteins will have a much higher abundance from the heavy sample.

To identify new Cyr1-binding proteins, we constructed a *C. albicans* strain for efficient labeling with stable isotopes. We then expressed a Myc-tagged Cyr1 in this strain to perform SILAC-based quantitative proteomics. We identified 36 novel Cyr1-interacting proteins and validated and characterized the interaction with the most confident candidate by conventional methods. Our results establish a global protein interaction network of Cyr1 and suggest the involvement of cell wall proteins in the Cyr1-mediated cAMP signaling pathway.

## 2. Materials and Methods

### 2.1. Strains, plasmids, and growth conditions

Yeast strains and plasmids used in this study are described in Table 1 and 2, respectively. Recombinant DNA manipulations were performed according to standard methods. *E. coli* XL1 blue (Stratagene, Cat. 200228) was used as the host strain for recombinant plasmids and cultured in LB broth (0.5% yeast extract, 1% tryptone, and 0.5% NaCl, pH 7.0) supplemented with 100 μg/ml ampicillin. Transformation of *C. albicans* was performed according to the protocol of the Fast Yeast Transformation Kit (G-Biosciences, Cat. GZ-1). Gene deletion was verified by colony PCR as described [24]. Looping out of *URA3* via FLP-mediated excision followed previous protocols [25]. Yeast cells were routinely grown at 30°C in YPD (2% yeast extract, 1% peptone, and 2% glucose) or GMM (glucose minimal medium, 6.79 g/l yeast nitrogen base without amino acids, and 2% glucose) supplemented with appropriate nutrients (80 μg/ml uridine, 40 μg/ml arginine, 40 μg/ml histidine, and 50 μg/ml lysine), and 1 mg/ml 5-FOA when necessary. Solid medium plates were prepared by adding 2% agar.

**Table 1.** *C. albicans* strains used in this study

| Strain | Relevant genotype |
| --- | --- |
| BWP17 | *ura3::imm434/ura3::imm434 his1::hisG/his1::hisG arg4::hisG/arg4::hisG* |
| GZY790 | BWP17, *lys2Δ::FRT/lys2Δ::FRT* |
| GZY808 | GZY790, *P<sub>MET3</sub>-Myc-URA3* |
| GZY809 | GZY790, *CYR1/cyr1::P<sub>MET3</sub>-Myc-CYR1-URA3* |
| GZY835 | BWP17, *CYR1/cyr1::P<sub>MET3</sub>-Myc-CYR1-URA3* |
| GZY865 | BWP17, *MP65/mp65::MP65-HA-HIS1* |
| GZY866 | BWP17, *CYR1/cyr1::P<sub>MET3</sub>-Myc-CYR1-URA3 MP65/mp65::MP65-HA-HIS1* |
| GZY871 | BWP17, *mp65Δ::UFP/mp65Δ::HIS1* |
| GZY888 | BWP17, *mp65Δ::FRT/mp65Δ::HIS1* |
| GZY900 | BWP17, *mp65Δ::FRT/mp65Δ::HIS1 P<sub>MET3</sub>-MP65-GFP-URA3* |
| GZY1094 | BWP17, *bcr1Δ::UFP/bcr1Δ::HIS1* |

**Table 2.**
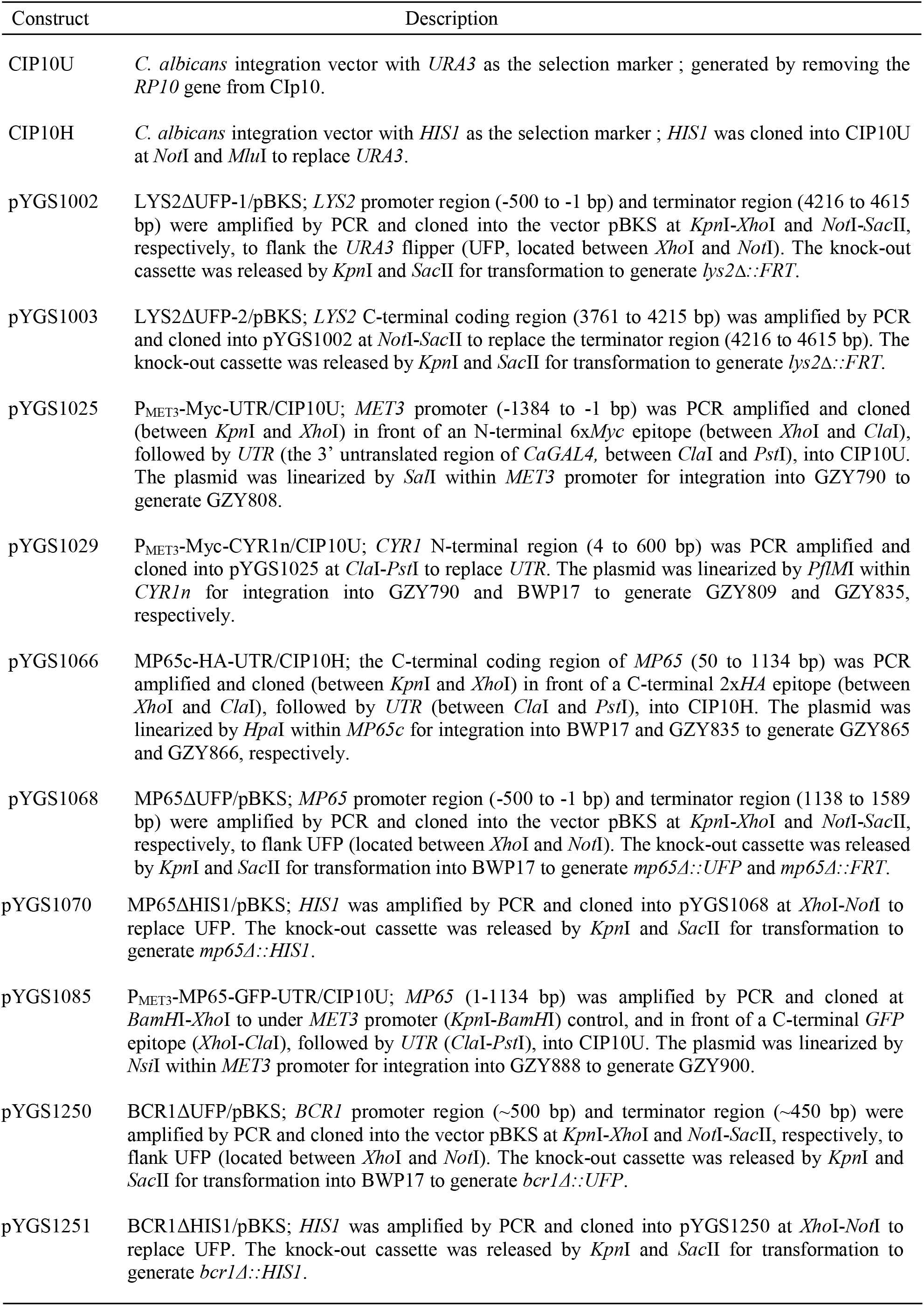
Plasmid constructs used in this study

For filamentous growth on Spider plates (10 g/l nutrient broth, 10 g/l mannitol, 2 g/l k2HPO4, and 20 g/l agar, pH 7.2), yeast cells were inoculated into YPD and cultured at 30°C overnight. Cells were then streaked onto plates to form single colonies by incubation at 30°C for 6 d. Images of filaments along a colony edge were taken using a Leica LEITZ DM RB optical microscope equipped with a Moticam 10MP digital camera. For hyphal induction, the YPD culture was 1:10 diluted into fresh YPD containing 10% fetal bovine serum (HyClone, Cat. SH30070.03) and incubated at 37°C for 2 h. Cells were visualized with a Leica DMRXA2 microscope equipped with a Coolsnap HQ2 digital camera. Images were acquired using MetaMorph 7.5 software.

For the Congo Red sensitivity test, yeast cells were grown to the stationary phase in YPD at 3Ü°C for 2 d. The cell suspensions were then 1:10 serially-diluted, spotted onto YPD plates containing 100 μg/ml of Congo red (Sigma-Aldrich, Cat. C6277-25G), and incubated at 3Ü°C for 24 h.

### 2.2. Protein extraction, immunoprecipitation (IP), and Western blotting (WB)

To prepare protein extracts, cells were harvested into 2 ml screw-cap microcentrifuge tubes by brief centrifugation to obtain pellets with a volume ≤ 500 μl and resuspended in an equal volume of ice-cold lysis buffer (50 mM Tris-HCl [pH7.4], 150 mM KCl, 1% NP-40) containing the protease inhibitor cocktail (Nacalai Tesque, Cat. 03969-21) and phosphatase inhibitor cocktail B (Santa Cruz, Cat. sc-45045). After adding an equal volume of acid-washed glass beads (Sigma-Aldrich, Cat. G8772-1KG), cells were broken by 6 rounds of 60-second beating at 5000 rpm in a TOMY Micro Smash beater (MS-100) with 1 min of cooling on ice between rounds. The lysed cells were then centrifuged at 14,000 rpm for 15 min at 4°C to collect the supernatant.

To perform IP, cell lysates were incubated with 30 μl slurry of goat polyclonal HA (Santa Cruz, Cat. sc-805 AC) or Myc (Santa Cruz, Cat. sc-789 AC) beads at 4°C for ≥ 1 h. After brief centrifugation at 8000 rpm, the beads were washed by resuspension in 800 μl cold lysis buffer, followed by another brief centrifugation. The washed beads were resuspended in 10 μl 1× protein loading buffer (50 mM Tris-HCl [pH6.8], 2% SDS, 10% glycerol, 1%β-mercaptoethanol, and 0.02 % bromophenol blue) and boiled for 6 min. Protein samples were separated by SDS-PAGE and transferred to a PVDF membrane (Bio-Rad, Cat. 162-0177).

For WB analysis, the membrane was first incubated in 5% milk in phosphate-buffered saline containing 0.1% Tween-20 (PBST) at room temperature for 1 h or at 4°C overnight. After a brief rinse with PBST, the membrane was incubated in PBST containing a 1:1000 diluted mouse monoclonal HA (Santa Cruz, Cat. sc-57594) or Myc (Santa Cruz, Cat. sc-40) antibody at room temperature for 1 h, followed by 3 rounds of 5-min wash with PBST. The membrane was then incubated in PBST containing a 1:2500 diluted secondary antibody (horseradish peroxidase-linked anti-mouse IgG from sheep; GE Healthcare, Cat. NA931V). After 3 rounds of 5-min wash with PBST, the membrane was immersed in ECL western blotting substrate solution (Thermo Scientific, Cat. 32209) and exposed to X-film (Fijifilm, Cat. 47410 19289).

### 2.3. Sample preparation for SILAC incorporation assay

GZY790 cells were inoculated into 2 ml of GMM supplemented with histidine, uridine, arginine-d10 (Cambridge Isotopes, Cat. CNLM-539-0.5), and lysine-d8 (Cambridge Isotopes, Cat. CNLM-291-0.5) (heavy medium) and grown at 30°C overnight. The culture was re-inoculated into 20 ml of the heavy medium and grown at 30°C for one more day. Cells were then harvested and protein extract was prepared with urea lysis buffer (20 mM HEPES [pH 8.0], 9 M urea, 1 mM sodium orthovanadate, 2.5 mM sodium pyrophosphate, and 1 mM ß-glycerophosphate). Mass spectrometry (MS) samples were prepared using the Filter-aided sample preparation (FASP) protocol [26]with some modifications. Briefly, 1:10 (v/v) of 50 mM dithiothreitol (DTT) was added to the lysate (100 μg) and incubated at 60°C for 20 min with shaking. 8M urea in 0.1 M Tris/HCl [pH 8.5] (UA) was added to the lysate and transferred to a Millipore 30 kDa Amicon filter unit. Buffer exchange was done on this filter unit with UA before alkylation with 50 mM iodoacetamide (IAA) in the dark for 20 min. Trypsin (porcine, modified sequencing grade; Promega) digestion (1:100) in 40 mM ammonium bicarbonate (ABC, Sigma-Aldrich) was performed on the filter at 37°C overnight. The next day, peptides were eluted by centrifuging at 20,000× g for 10 min at room temperature, with additional elution using 40 mM ABC. Total tryptic digested peptides were concentrated using a speed vacuum concentrator before introducing them into the mass spectrometer for incorporation analysis.

### 2.4. Immuno-purification of Myc-Cyr1-associated proteins

The strain expressing Myc-Cyr1 (GZY809) was inoculated into 50 ml of the heavy medium and grown at 30°C overnight. The culture was reinoculated into 500 ml of the heavy medium and grown at 30°C for one more day. The control strain GZY808 was cultured similarly in the light medium (GMM supplemented with histidine, uridine, arginine, and lysine). Cells were then harvested into 2 ml screw-cap microcentrifuge tubes, and 7.5 ml of heavy and light protein extracts were prepared and incubated with 40 μl EZview red anti-Myc affinity gel (Sigma-Aldrich, Cat. E6654-1ML) at 4°C for 4 h. After washing with 1 ml of lysis buffer 2 times, the beads were combined and washed for 3 more times, and finally resuspended in 40 μl of 1× protein loading buffer and boiled for 6 min. Eluted protein complexes were separated by one-dimensional 4-12% NuPage Novex Bis-Tris Gel (Invitrogen, Cat. NP0321BOX) and stained using the Colloidal Blue Staining Kit (Invitrogen, Cat. LC6025). All visualized protein bands were excised for quantitative proteomic analysis.

### 2.5. Quantitative mass spectrometry (MS)

SDS-PAGE-separated proteins were digested in-gel with trypsin using published protocols [27]. Peptide samples were analyzed on an Orbitrap (Thermo Fisher). Survey full-scan MS spectra (m/z 310–1400) were acquired with a resolution of r = 60,000 at m/z 400, an AGC target of 1e6, and a maximum injection time of 500 ms. The ten most intense peptide ions in each survey scan with an ion intensity of >2000 counts and a charge state ≥2 were isolated sequentially to a target value of 1e4 and fragmented in the linear ion trap by collision-induced dissociation using a normalized collision energy of 35%. Dynamic exclusion was applied using a maximum exclusion list of 500 with one repeat count, repeat, and exclusion duration of 30 s. Raw data were searched using C_albicans_SC5314_version _A21-s02-m01-r02_orf_trans_all.fasta. Database searches were performed with tryptic specificity allowing maximum of two missed cleavages and two labeled amino acids and an initial mass tolerance of 6ppm for precursor ions and 0.5Da for fragment ions. Cysteine carbamidomethylation was searched as a fixed modification, and N-acetylation and oxidized methionine were searched as variable modifications. Labeled arginine and lysine were specified as fixed or variable modifications, depending on the prior knowledge about the parent ion. SILAC peptide and protein quantification was performed with MaxQuant version 1.2.0.18 using default settings. Maximum false discovery rates were set to 0.01 for both protein and peptide. Proteins were considered identified when supported by at least one unique peptide with a minimum length of six amino acids.

### 2.6. Biofilm development and quantification

Biofilm development assay was performed using the method described previously with slight modifications [28]. Single colonies were inoculated in liquid GMM and incubated at 30°C overnight with shaking at 180 rpm. Cells were washed twice with PBS. The cell concentration was adjusted to 10^7^ cells/ml PBS using a haemocytometer. 100 μl of the cell suspension was transferred to a well of a flat-bottomed 96-well polystyrene plate (Tissue culture-treated, Falcon) and incubated for 1.5 h at 37°C with shaking at 75 rpm to allow cell adherence. The plate was sealed with the Breathe-Easy sealing membrane (Axygen) to prevent cross-contamination between wells and edge effect. Next, the non-adherent cells were removed, and the well was washed twice with 150 μl of PBS. 200 μl of fresh GMM was added and incubated for 48 h at 37°C with shaking at 75 rpm. Media was changed after 24 h to remove planktonic cells. After 48 h incubation, planktonic cells were aspirated from the wells and the biofilms were washed twice with PBS prior to quantification via OD_600_ and metabolic activity measurement.

For biofilm visualization, the 96-well polystyrene plate was mounted on the stage of an inverted microscope after drying the biofilm. High-magnification images of *C. albicans* cells were taken with an Olympus IX70 microscope (Olympus, Tokyo, Japan) equipped with a Motic Cam 2.0 MP camera and Motic Images Plus 2.0 software.

For biofilm measurement, OD_600_ was read using a microplate reader (Tecan Infinite M200 Pro). Five independent locations per well were read to gain an averaged density. The metabolic activity of the biofilms was measured using the 2,3-bis-(2-methoxy-4-nitro-5-sulfophenyl)-2H-tetrazolium-5-carboxanilide, disodium salt (XTT) colorimetric assay (Biotium) according to the manufacturer’s protocol. 150 μl of PBS and 25 μl of activated XTT solution (Activation Reagent:XTT solution = 1:200) was added in each well. The plate was then incubated at 37°C for 2 h with shaking in the dark. 100 μl of the supernatant was transferred to a new well. The absorbance was read at 490 nm using a microplate reader. Graph Pad Prism Software Version 6.00 was used for all statistical analyses. Data were expressed as ±standard deviations. Results of biofilm development assay were analyzed by two-tailed unpaired *t*-test. All experiments were repeated at least three times independently with similar results.

## 3. Results

### 3.1. Construction of a C. albicans strain for SILAC-based proteomics

To achieve optimal separation between light and heavy peptides during quantitative proteomic analysis, we decided to label the proteome of *C. albicans* with two heavy amino acids, arginine-d10 (deuterated arginine) and lysine-d8 (deuterated lysine), simultaneously. To generate a strain auxotrophic for both arginine and lysine to allow metabolic labeling, we deleted the two alleles of the *LYS2* gene sequentially (Figure 1A) in the host strain BWP17 (ura^-^ his^-^ arg^-^) using a recyclable selection marker (the *URA3* flipper [25]), yielding the strain GZY790 (ura^-^ his^-^ lys^-^ arg^-^). The auxotrophy of GZY790 was confirmed as the strain failed to grow in the absence of any of the four nutrients arginine, histidine, lysine, and uridine (Figure 1B).

**Figure 1.**
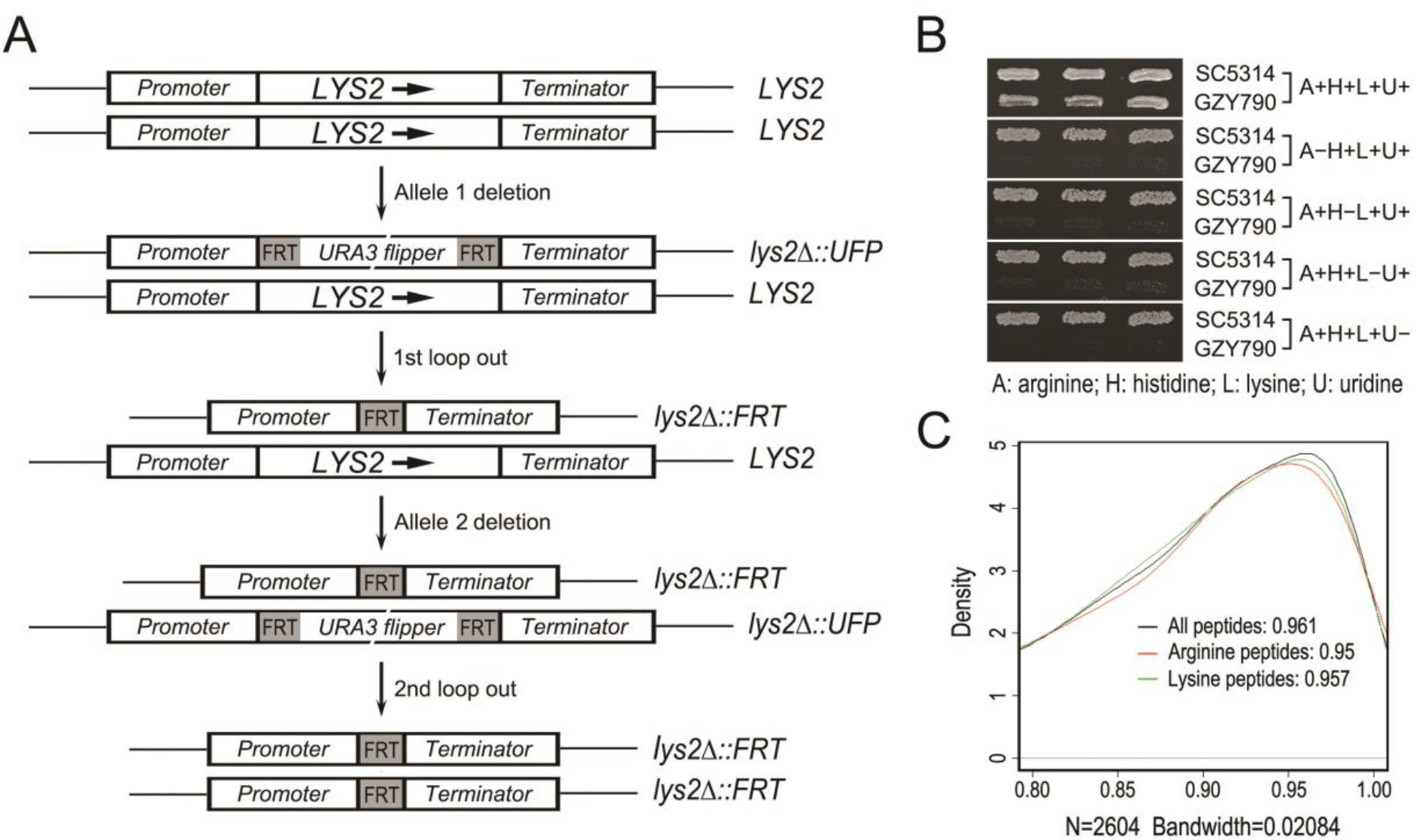
Construction of a strain optimized for SILAC experiments. (**A**) A schematic diagram showing the steps to delete the *LYS2* gene in BPW17 to generate GZY790. (**B**) Confirmation of the auxotrophy of GZY790. Cells of GZY790 and the WT strain SC5314 were grown on GMM plates supplemented with arginine, histidine, lysine, and uridine, and replicated to GMM plates lacking one of the four nutrients as indicated. Plates were incubated at 30°C overnight. (**C**) Incorporation rate assay of GZY790. The strain was grown in the GMM medium supplemented with histidine, uridine, and heavy isotope-labelled arginine-d10 and lysine-d8. Cells were then harvested and protein extracts prepared to assess the incorporation rate.

Many factors, such as protein synthesis and degradation, cell division, and growth conditions, may affect the incorporation rate of heavy amino acids into the proteome of a strain. To assess the efficiency of metabolic labeling, GZY790 was cultured in heavy Glucose Minimal Medium (GMM), in which arginine and lysine were provided in the form of arginine-d10 and lysine-d8, at 30°C for 40 h. Protein extracts were then prepared to determine the percentage of incorporation of the heavy amino acids. We found that 95% of peptides were successfully labeled with arginine-d10, while 95.7% were labeled with lysine-d8 (Figure 1C). When peptides labeled with either arginine-d10 or lysine-d8 or both were counted, the percentage increased to 96.1% (Figure 1C). Such a high incorporation rate met the requirement of SILAC-based quantitative proteomic studies. However, when 10-20% serum was added to the medium in order to induce hyphal growth, the incorporation rate of heavy amino acids was reduced to lower than 90%, most likely owing to the presence of natural amino acids in serum, which is not suitable for the SILAC-based analysis. Thus, our studies described below were carried out only under yeast growth conditions by using GMM without serum.

### 3.2. Purification of Cyr1-associated proteins for SILAC MS

To identify Cyr1-associated proteins by SILAC-based quantitative proteomics, we tagged Cyr1 with an N-terminal Myc epitope under the control of the *MET3* promoter in GZY790. The expression of Myc-tagged Cyr1 in the resulting strain GZY809 was verified by immunoprecipitation (IP) and Western blotting (WB) with anti-Myc (α-Myc) antibodies (Figure 2A).

**Figure 2.**
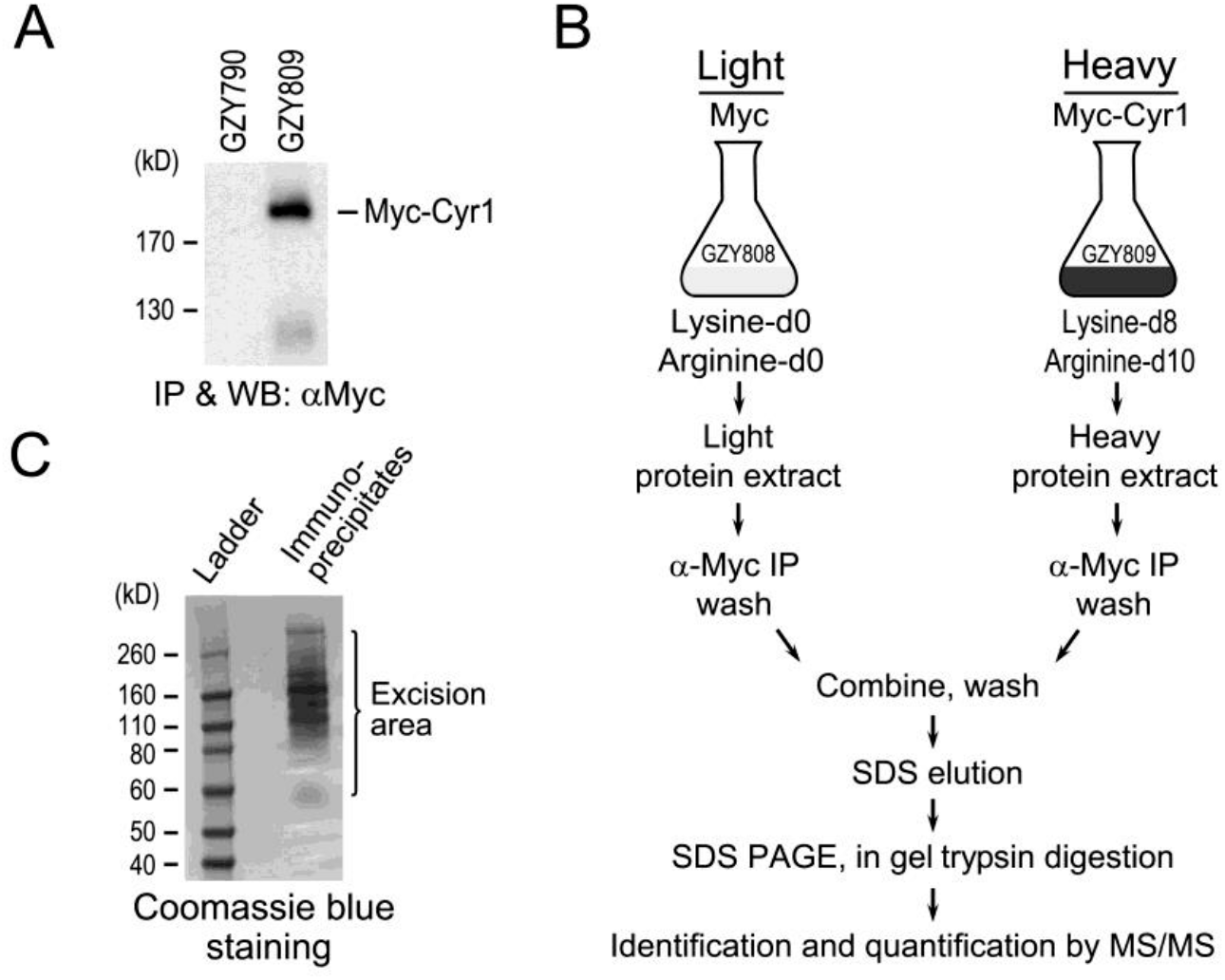
Pull down of Myc-Cyr1-associated proteins for quantitative analysis. (**A**) Tagging of Cyr1 with an N-terminal Myc epitope in GZY790. Anti-Myc IP and WB confirmed the expression of Myc-Cyr1 in the resulting strain (GZY809). (**B**) Experimental workflow used for the identification and quantification of Myc-Cyr1-associated proteins. GZY809 and the control strain GZY808 were grown in heavy and light media. Cells were harvested and protein extracts prepared for IP with a Myc antibody conjugated on agarose beads. The beads were combined in a 1:1 ratio during washes. Beads-bound proteins were eluted with SDS and subsequently separated by SDS-PAGE and digested with trypsin for quantitative analysis. (**C**) Visualization of Myc-Cyr1 associated proteins. The combined immunoprecipitates from GZY808 and GZY809 were separated by SDS-PAGE and stained with Coomassie blue. All visible bands were excised for in-gel trypsin digestion.

The procedures to quantitatively characterize Cyr1-interacting partners are schematically illustrated in Figure 2B (see Materials and Methods for details). To metabolically label the whole proteome of GZY809, cells of the strain were grown in the heavy medium for 40 h to allow maximum incorporation of heavy amino acids. For control, a strain expressing only the Myc epitope (GZY808) was cultured in the light medium under otherwise the same conditions. Equal amounts of heavy and light protein extracts were prepared and incubated with agarose beads conjugated with the Myc antibody separately. The beads from the two samples were then combined for washes. Finally, the beads-bound proteins were eluted with SDS, separated by SDS-PAGE, and visualized with Coomassie blue staining (Figure 2C). All proteins bands were excised, subjected to in-gel trypsin digestion, and analyzed using a nanoLC Orbitrap MS system.

### 3.3. Identification of Cyr1-binding partners by quantitative analysis

A total of 670 proteins were identified from the immunoprecipitates of Myc-Cyr1 (Table S1). SILAC MS made it possible to compare the amounts of proteins immunoprecipitated from cells expressing Myc (GZY808, light) or Myc-Cyr1 (GZY809, heavy). The integrated intensity sum of the peptide peaks as determined by MaxQuant reflects the peptide abundance, and the fold change represents the ratio of peptides quantified between the experimental sample labeled with heavy amino acids and the control sample labeled with light amino acids. A plot of “Ratio vs Log10(Intensity)” was then drawn to display all the identified proteins as dots in a map; and the identity of the proteins with the highest possibility to be Cyr1 interactors were labeled as yellow and red dots (Figure 3A). Many proteins identified by SILAC MS exhibited a normalized heavy:light (H/L) ratio below 1:1 (Figure 3B), and therefore likely to represent the background and/or non-specific protein interactions. The greater the peptide H/L ratio’s deviation is from 1:1, the higher the likelihood they are true Cyr1-binding partners. By using a H/L ratio of 1.5:1, we identified 38 high-priority Cyr1 interacting proteins (Figure 3C), including two previously characterized Cyr1-associated proteins, Cap1 (normalized H/L ratio of 59) and Act1 (normalized H/L ratio of 2.7) [22].

**Figure 3.**
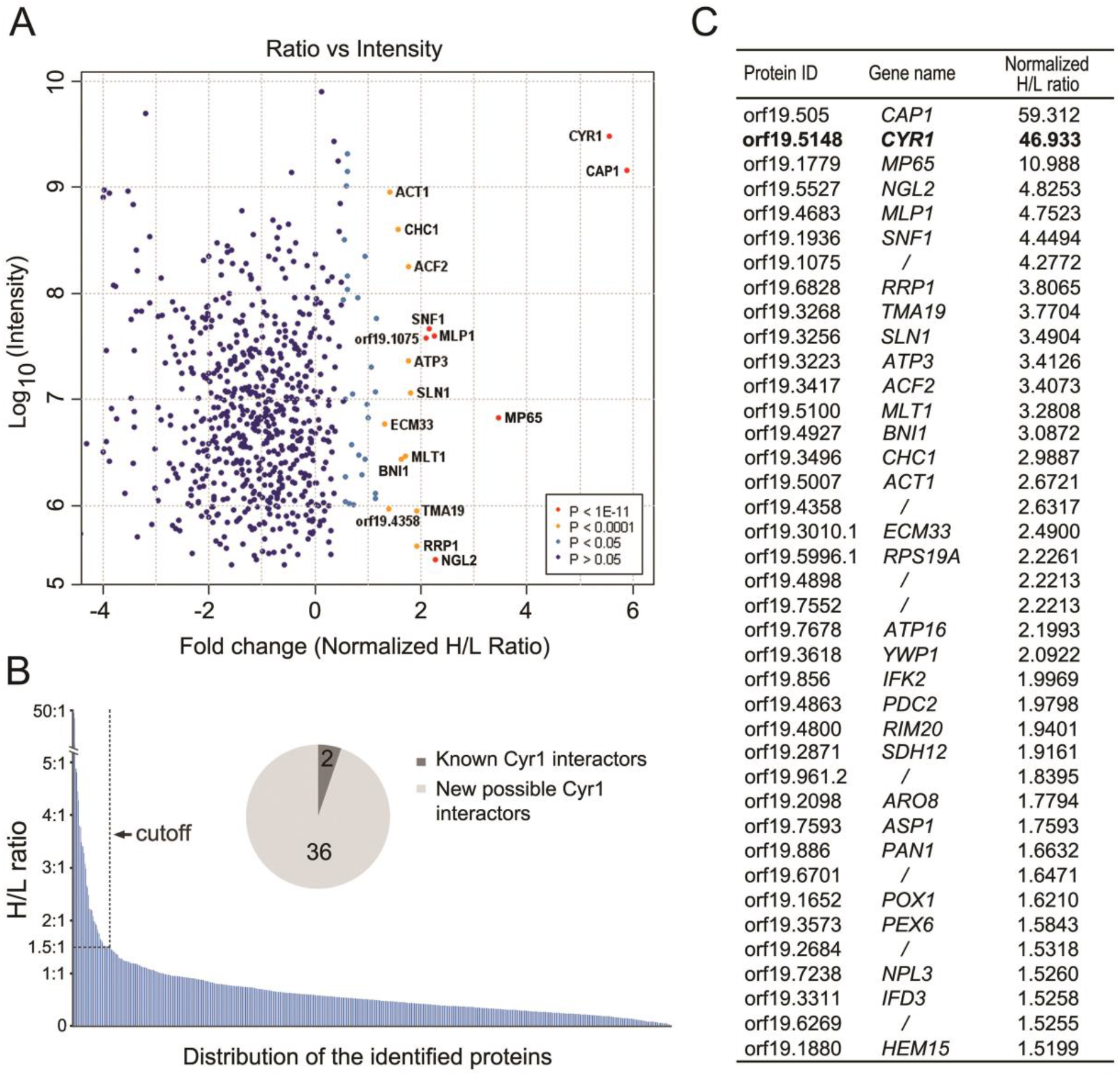
Quantitative analysis of Myc-Cyr1 associated proteins. (**A**) Ratio vs intensity plot of identified proteins. The trypsin-digested immunoprecipitates from GZY808 and GZY809 were processed for protein identification and quantitation. Each protein’s heavy:light ratio was plotted against its total intensity (in Log10 value). Red and yellow dots (labeled with protein ID) indicate the potential interactors of Cyr1. (**B**) Distribution of the identified proteins according to their normalized heavy:light ratios. Proteins associated with a heavy:light ratio of more than 1.5:1 are potential Cyr1 interactors. The pie chart represents the proportion of previously known and newly identified Cyr1-interacting proteins. (**C**) List of the protein identities of potential Cyr1 interactors with the corresponding gene names and normalized heavy:light ratios. The bait protein Cyr1 is highlighted in bold.

### 3.4. Characterization of Mp65

Identifying Cap1 and Act1 among the top potential Cyr1 interactors provides strong validation for the SILAC-based method. Among the remaining 36 highly confident candidates, Mp65 has the highest normalized H/L ratio of ~11 (Figure 3C). This protein is a putative β-glucanase mannoprotein and has been previously reported to have a role in several virulence-related traits such as cell wall integrity, adhesion, biofilm formation, hyphal morphogenesis and pathogenicity of *C. albicans* [29,30]. Therefore, we selected Mp65 for further investigation.

To verify that the Mp65 protein is an authentic Cyr1-binding partner, we tagged Mp65 with a C-terminal HA epitope and co-expressed it with Myc-Cyr1 for co-IP experiments. When the HA antibody was used to pull down Mp65-HA from cell lysates, Myc-Cyr1 was detected in the IP product by anti-Myc WB (Figure 4A, left). Reciprocally, when Myc-Cyr1 was precipitated with the Myc antibody, Mp65-HA was detected by anti-HA WB (Figure 4A, right). In the control experiments, the HA antibody failed to pull down Myc-Cyr1 from cells expressing Myc-Cyr1 alone, and the Myc antibody failed to pull down Mp65-HA from cells where Cyr1 was untagged (Figure 4A). Taken together, these experiments demonstrate that Cyr1 and Mp65 physically interact with each other *in vivo*.

**Figure 4.**
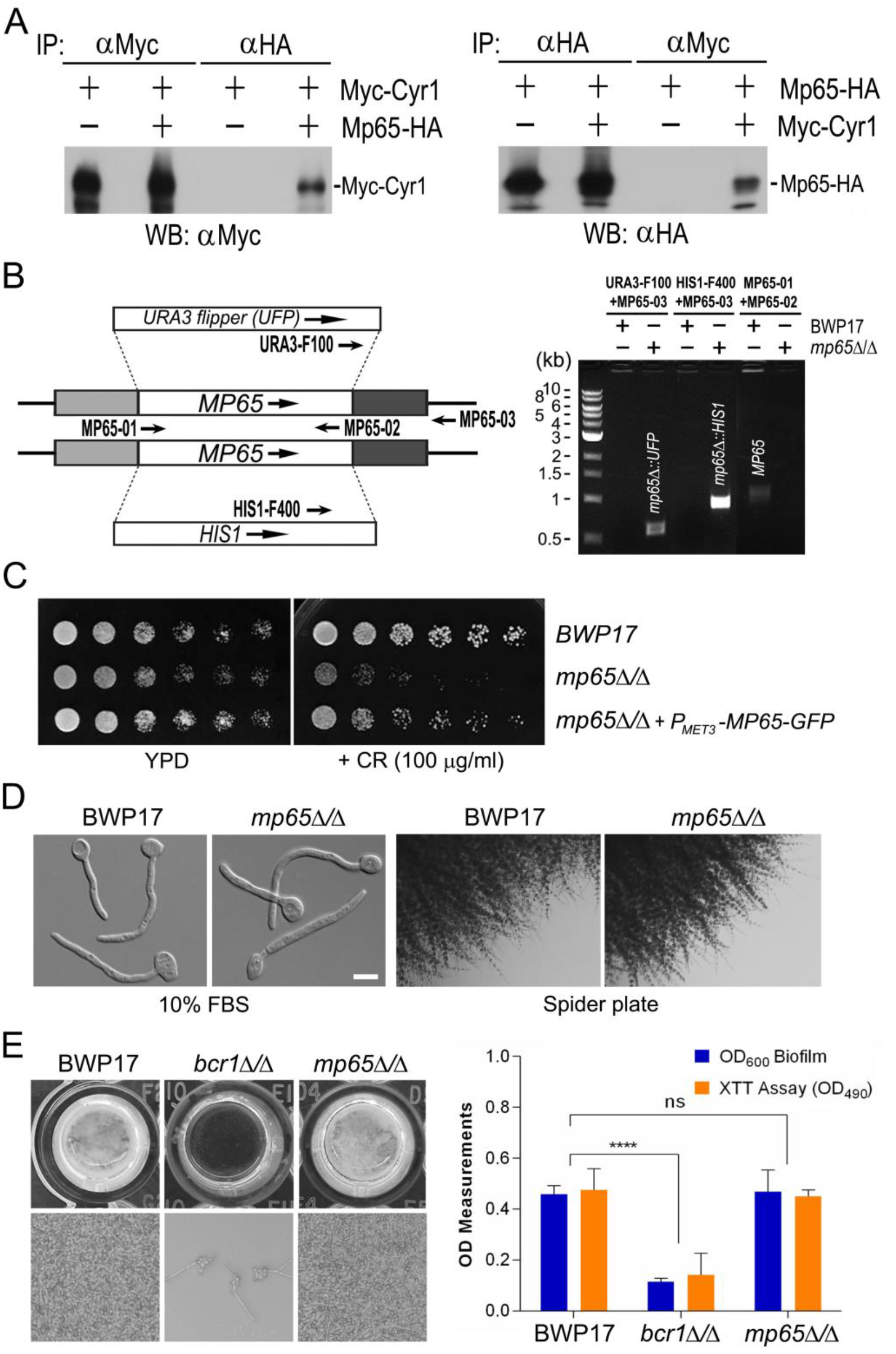
Characterization of Mp65. (**A**) Validation of the physical interaction between Cyr1 and Mp65 by co-IP. Protein extracts prepared from GZY835 (Myc-Cyr1) and GZY866 (Myc-Cyr1 Mp65-HA) were subjected to either anti-HA or anti-Myc IP and probed with the Myc antibody (left). Similarly, protein extracts prepared from GZY865 (Mp65-HA) and GZY866 (Myc-Cyr1 Mp65-HA) were subjected to either anti-HA or anti-Myc IP and probed with the HA antibody (right). (**B**) Deletion of the *MP65* gene in BWP17. The *UFP* and *HIS1* selectable marker genes were used to replace the two alleles of *MP65*. Colony PCR was carried out with primers URA3-F100 (caatcaaaggtggtccttctgcag) and MP65-03 (tcaaaacaatgac ttcacatat) to verify the correct *UFP* replacement of *MP65*, and with primers HIS1-F400 (acgaattgaagaaagctggtgc) and MP65-03 to verify the *HIS1* replacement of *MP65*. The primers MP65-01 (tgctccattagctcatcaacatca) and MP65-02 (ttagagtaaataccccagtatttt) were used for colony PCR to confirm the absence of *MP65* gene in the *mp65*Δ/Δ mutant (GZY871). (**C**) Congo Red (CR) sensitivity test of the *mp65*Δ/Δ mutant. YPD cultures of BWP17, GZY871 (*mp65*Δ/Δ), and GZY900 (*mp65*Δ/Δ::*P_MET3_-GFP-MP65*) were serially diluted (1:10) in water and spotted onto YPD plates without or with 100 μg/ml of CR. The plates were incubated at 30°C for 24 h. (**D**) Filamentous growth of the *mp65*Δ/Δ mutant. For hyphal morphology observation, YPD cultures of BWP17 and GZY871 (*mp65*Δ/Δ) were induced for hyphal growth in the YPD medium containing 10% fetal bovine serum (FBS) at 37°C for 2 h. For filamentous growth on Spider plates, YPD cultures of BWP17 and GZY871 (*mp65*Δ/Δ) were streaked onto Spider plates and incubated at 30°C for 6 d. (**E**) Biofilm development assays of the *mp65*Δ/Δ mutant. Biofilms of BWP17, GZY1094 (*bcrl*Δ/Δ), and GZY871 (*mp65*Δ/Δ) were formed on the bottom of a 96-well polystyrene plate and subjected to visual inspection and microscopic examination. Biofilms for each strain were further quantified by OD_600_ biofilm measurement and XTT assay at OD490. BWP17 was used as the positive control (normal biofilm) and *bcr1*Δ/Δ was used as negative control (biofilm defects). \*\*\*\**P* <0.0001. *mp65*Δ/Δ displayed no significant (ns) differences in OD_600_ (*P*=0.7441) and XTT assay (*P*=0.4147) as compared to BWP17.

To explore the functional relationship between Mp65 and Cyr1, we generated an *mp65*Δ/Δ mutant (GZY888) using the common laboratory tool strain BWP17. Both wild-type (WT) copies of *MP65* were replaced by the auxotrophic markers *UFP* and *HIS1*, respectively, and the deletion of *MP65* was verified by colony PCR (Figure 4B) according to standard protocols [24]. Consistent with a previous study [29], our *mp65*Δ/Δ mutant was sensitive to the cell wall perturbing agent Congo Red (CR) (Figure 4C). When the *mp65*Δ/Δ culture was serially diluted and spotted onto YPD plates, the cells grew well in the absence of CR, but failed to grow on plates containing 100 μg/ml of CR. In comparison, BWP17 cells grew normally in the presence of 100 μg/ml of CR. When a WT copy of *MP65* was reintroduced into the mutant, the CR sensitivity was completely rescued, indicating that the cell wall defect of *mp65*Δ/Δ was caused by the loss of *MP65*. These results confirm that *MP65* is required for cell wall integrity,

Unexpectedly, in contrast to the previous report [30], our *mp65*Δ/Δ mutant exhibited normal hyphal growth like BWP17 (Figure 4D). When cultured in YPD containing 10% serum at 37°C for 2 h, the *mp65*Δ/Δ cells produced thin, long hyphae morphologically indistinguishable from BWP17. Moreover, the *mp65*Δ/Δ mutant also underwent normal filamentous growth on the solid medium (Figure 4D). After 6 d incubation at 30°C on Spider plates, filaments with short branches of comparable lengths were seen along the edges of both *mp65*Δ/Δ and BWP17 colonies. In addition, our *mp65*Δ/Δ mutant exhibited normal biofilm formation like its parental WT strain BWP17 (Figure 4E). After incubation in the GMM medium at 37°C for 48 h in a flat-bottomed 96-well polystyrene plate, both strains formed healthy biofilms with compatible architecture and quantity, which were in sharp contrast with the biofilm-defective strain *bcr1*Δ/Δ [31] (Figure 4E).

### 3.5. Classification of the identified Cyr1-binding partners

To better interpret the proteomics data, we grouped the potential Cyr1-interacting proteins, including the two known interacting partners, Cap1 and Act1, based on their molecular functions and biochemical properties but with an emphasis on established protein-protein interactions (Figure 5).

**Figure 5.**
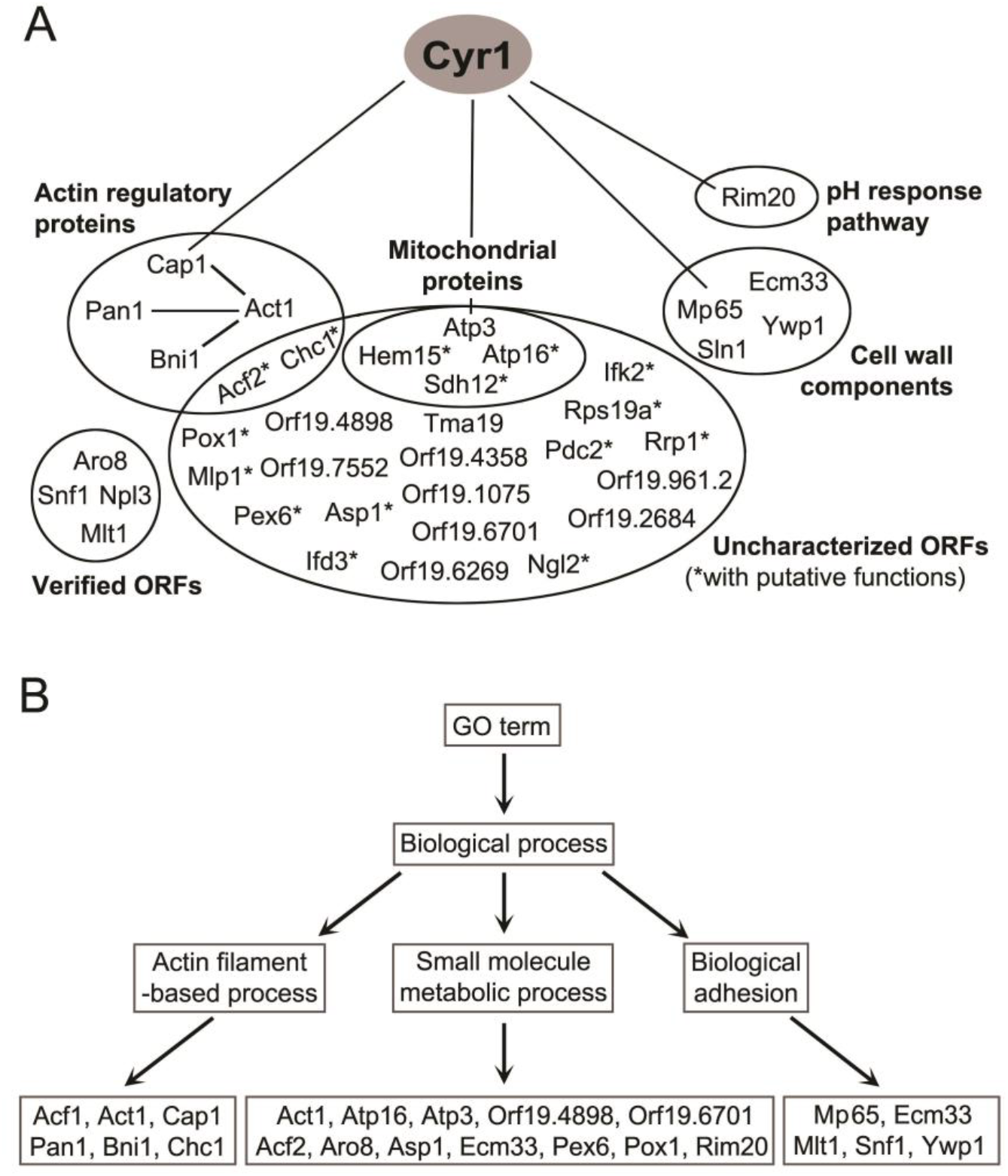
Classification of potential Cyr1 interactors identified by the SILAC-based approach. (**A**) The 36 potential Cyr1 interactors with two known binding partners (Cap1 and Act1) were manually curated into different groups with some members overlapped. (**B**) Query of the Cyr1 potential interactors with Gene Ontology (GO) Term Finder (*Candida* Genome Database) under “Biological process” ontology classified some of the candidates into three categories: “actin filament-based process”, “small molecule metabolic process”, and “biological adhesion”.

When we analyzed the molecular functions of the potential Cyr1 interactors using Gene Ontology Term Finder from the *Candida* Genome Database, no significant ontology term can be found for input genes. Therefore, we turned to manual curation (Figure 5A). Among the 38 identified proteins, more than half of the candidates are uncharacterized and therefore classified into a group of “uncharacterized ORFs”. In this group, some proteins have putative functions inferred from their characterized orthologues in other species (marked with an asterisk). Four proteins within this group (Atp3, Atp16, Hem15, and Sdh12) have mitochondrion-related functions and form a subgroup of “mitochondrial proteins”. Four actin-related proteins, Bni1, Pan1, Chc1, and Acf2, together with Cap1 and Act1, form a “actin regulatory proteins” group. These proteins either bind to actin or regulate cortical actin patch dynamics [22,32–36]. We classified Mp65 with Sln1, Ywp1 and Ecm33 into a “cell wall components” group, as these proteins either localize to the cell wall and/or regulate cell wall biosynthesis [29,37–40]. Rim20 is the only potential Cyr1 interacting partner that functions in the pH response pathway [41]. Four other verified ORFs (Aro8, Snf1, Npl3, and Mlt1) do not belong to any of the above groups and thus are classified as “verified ORFs”.

To identify possible biological processes in which the potential Cyr1-binding partners are involved, we also input the 38 candidates to Gene Ontology Term Finder and performed a search using the default setting. In addition to the “small molecule metabolic process”, two prominent biological processes were identified: one is “actin filament-based process” which consists of Act1, Bni1, Cap1, Pan1, Chc1, and Acf2, and the other is “biological adhesion” that contains Mp65, Ecm33, Ywp1, Mlt1, and Snf1 (Figure 5B). These findings are largely consistent with our own classification above.

## 4. Discussion

As the sole adenylyl cyclase of the human fungal pathogen *C. albicans*, Cyr1 catalyzes cAMP synthesis in response to many external and internal stimuli and triggers the cAMP-dependent signaling pathway to initiate proper physiological responses. Although several factors that regulate the catalytic activity of Cyr1 have been characterized, how Cyr1 senses and integrates the wide range of external and internal signals with distinct nature remains largely unknown. In this work, we applied SILAC-based approaches to analyze the interactome of Cyr1 comprehensively. Our study has identified 36 novel candidates as potential interactors of Cyr1, thus providing a valuable resource for discovering novel molecular mechanisms that regulate Cyr1 activities in response to various stimuli.

### 4.1. The SILAC-optimized C. albicans strain GZY790

Since its development in 2002, SILAC-based quantitative proteomic analysis has become an excellent popular technique to reveal changes in protein abundance and identify protein interaction partners in many organisms. The metabolic labeling of SILAC has proven to have unique and specific advantages compared to other labeling methods. However, SILAC has limitations in its application because human tissue/organ samples and some types of unicellular organisms may encounter difficulties in incorporating heavy amino acids. Until recently, the utilization of SILAC to *C. albicans* has been proven applicable and used to identify numerous interactors of the protein phosphatase Cdc14 [42,43].

For a successful SILAC experiment, the choice of heavy-isotope-labeled amino acids is crucial. The heavy amino acids are required to provide at least 4-Da separation of heavy and light peptides to reduce the overlapping of light and heavy peptide clusters and inaccuracy in quantitation. The commonly used laboratory strain of *C. albicans*, BWP17, is only suitable for metabolic labeling with heavy arginine since it is auxotrophic for uridine, histidine, and arginine [44]. To ensure better resolution of heavy and light peptide pairs, we further introduced lysine auxotrophy into BWP17 by deleting the *LYS2* gene to generate the strain GZY790. Being auxotrophic for both arginine and lysine, GZY790 allows the incorporation of heavy arginine and lysine into *C. albicans* proteome simultaneously at a rate higher than 95%, rendering it a useful tool strain for SILAC-based quantitative proteomics. The identification of 36 new Cyr1-interacting partners together with 2 well-characterized Cry1-binding proteins by using GZY790 demonstrates the efficacy of this strain.

### 4.2 The identification of novel Cyr1-interacting proteins

SILAC-based metabolic labeling in conjugation with MS analysis enabled us to reveal the identity of the proteins co-immunoprecipitated with Myc-Cyr1, and at the same time, exclude the nonspecific binding proteins by quantitative analysis. The nonspecific binding proteins were most likely pulled down during IP due to their association with the Myc epitope, the Myc antibody, and/or the antibody-conjugated beads, rather than with Cyr1 itself, thus showing a similar abundance in light and heavy samples. It is expected that the SILAC-MS method will identify genuine Cyr1-interacting proteins. Indeed, we identified Cap1 and Act1 with high confidence, two proteins previously shown to form a stable complex with Cyr1 [22]. We also discovered 36 more proteins as novel Cyr1 interacting proteins, revealing a comprehensive interactome for Cyr1. One of the novel proteins, Snf1, has been confirmed to physically interact with Cyr1 in *S. cerevisiae* [45]. However, we failed to detect some other proteins shown or thought to interact with Cyr1 by other methods, such as the small GTPase Ras1 [17]. Two possible explanations are: first, the interaction between Ras1 and Cyr1 requires activation by external stimuli such as serum, while our experiments were done in yeast cells. Second, the association of these two proteins is transient and therefore difficult to detect. This may also explain why other potential Cyr1-interacting proteins, Gpa2 and Hsp90/Sgt1, were not picked up in this experiment. The association of Gpa2 with Cyr1 may require the external inducers glucose and/or amino acids [15,16], while the association of Sgt1 with Cyr1 may happen only at elevated temperature [19]. Therefore, it is necessary to perform the same experiment under different growth conditions, such as filamentous growth induced by various stimuli, to construct a more comprehensive interactome for Cyr1.

### 4.3. Actin regulatory proteins as Cyr1 interactors

Among the identified novel Cyr1 interactors, several (Pan1, Bni1, Chc1, Acf2) are functionally related to actin dynamics. Together with Cap1 and Act1, these proteins form the group of actin regulatory proteins. Pan1 functions as an activator of the actin-nucleating Arp2/3 complex in *S. cerevisiae* [32]and has been shown to form a complex with the endocytic protein Sla1 to regulate cortical actin patch dynamics during the polarized growth of *C. albicans* [35]. Bni1 is the yeast formin that mediates actin filament assembly independent of the Arp2/3 complex [33]. In *C. albicans*, Bni1 plays an important role in cell polarity control during both yeast and hyphal growth [46]. Although the functions of Chc1 and Acf2 have not been characterized in *C. albicans*, the orthologue of Chc1 (clathrin heavy chain) in *S. cerevisiae* has been demonstrated to associate with the actin-based endocytic machinery [34]. The orthologue of Acf2 (assembly complementing factor) in *S. cerevisiae* is required for in vitro cortical actin assembly [36]. We noticed that two more actin-related proteins, Sla2 and Arc18, were also identified in the same experiment, but with a normalized H/L ratio (1.4153 and 1.177, respectively) slightly lower than the cutoff value we used (Supplemental Table 1). In *C. albicans*, Sla2 localizes to cortical actin patches and is required for the proper organization of the actin cytoskeleton and formation of true hyphae in response to serum and other stimuli [47,48]. The orthologue of Arc18 in *S. cerevisiae* is a subunit of the Arp2/3 complex that is required for the motility and integrity of cortical actin patches [49].

Because Cyr1 forms a stable tripartite complex with Act1 via Cap1 [22], it is not surprising that some actin-associated proteins were co-immunoprecipitated with Cyr1. However, we cannot rule out the possibility that some of the actin regulatory proteins may interact with Cyr1 directly, independent of Cap1 and Act1. What are the physiological roles for the interaction between Cyr1 and actin regulatory proteins? We previously proposed that the Cyr1/Cap1/actin complex may provide a mechanism that monitors the intracellular status of the actin cytoskeleton and adjust cAMP synthesis to best regulate actin-mediated cellular processes such as polarized growth [22,50]. Here, based on the discovery of more Cyr1-associated actin regulators, we expand our hypothesis that Cyr1 might directly interact with these proteins in organizing the actin cytoskeleton for polarized growth in response to external and internal stimuli. Because cortical actin patches are the active endocytic sites, it is also possible that these actin regulatory proteins may help to internalize some external inducers through the endocytic pathway and subsequently deliver the signals to Cyr1.

### 4.4. Mp65 and cell wall components as Cyr1 interactors

In addition to actin regulatory proteins, 4 proteins (Mp65, Ywp1, Ecm33, and Sln1) with cell wall-related functions have also been identified from the same experiment. Mp65 is a putative β–glucanase adhesin previously shown to be required for cell wall integrity and several infection-related cellular processes [29,30]. Ywp1 is a yeast cell wall protein that appears to be linked covalently to the glucan of the cell wall matrix [37]. Ecm33 is a GPI-anchored protein required for cell wall integrity, morphogenesis, and virulence of *C. albicans* [39]. Sln1 is a histidine kinase involved in a two-component signaling pathway regulating cell wall biosynthesis [38]. Together, these proteins define a novel group of Cyr1 interactors: cell wall components.

The cell surface localization of these cell wall proteins suggests a possible role in sensing and transmitting external signals to Cyr1. We detected Mp65 association with Cyr1 by both SILAC MS and co-IP, revealing a novel link between a surface protein and the intracellular Cyr1. Several highly possible transmembrane domains in Mp65 with regions exposed to both the outside and inside of the plasma membrane were identified by transmembrane topology analysis (http://www.ch.embnet.org/software/TMPRED_form.html) (Figure S1), providing potential binding sites for signaling molecules. It is tantalizing to propose that Mp65 transduces external signals across the plasma membrane to regulate Cyr1 activity. As several other cell wall proteins have also been found to interact with Cyr1, one or more of them may function in parallel with Mp65, which may explain why the deletion of *MP65* alone did not cause any apparent morphological defects. Since Mp65 has been demonstrated to be a major antigen target of cell-mediated immune response to *C. albicans* [51], it is reasonable to speculate that Mp65 (and possibly other cell wall proteins) may trigger the hyphal growth of *C. albicans* via the cAMP pathway to protect the pathogen against attacks by the human immune system. If this is true, Mp65 and other cell wall proteins identified in this study may provide a novel group of targets for both pharmacological and immunological antifungal therapies.

We confirmed that Mp65 is necessary for cell wall integrity, as previously reported [29]. However, we could not repeat the previous observations that the *mp65*Δ/Δ mutant was defective in hyphal morphogenesis and biofilm formation [30]. Our *mp65*Δ/Δ mutant produced filaments on solid Spider medium, formed normal hyphae in the serum-inducing medium, and developed robust biofilm in manners indistinguishable from WT strains. We are not sure what causes the discrepancy, although it is possibly due to variations in strain background and experimental conditions.

### 4.5. Rim20 as a Cyr1 interactor

pH is an important environmental variable that affects *C. albicans* morphogenesis, with alkaline conditions favoring hyphal growth and acid conditions favoring yeast growth [52]. *C. albicans* pH sensing and its morphological response to pH is primarily mediated by the Rim101 signal transduction pathway [41]. Recently, the cAMP/PKA signaling pathway is shown to be involved in the pH-dependent morphological responses of *C. albicans*. It was reported that acidic pH downregulated cAMP signaling and inhibited hyphal growth in minimal medium with GlcNAc. More importantly, such effect is thought to be mediated, at least in part, through Cyr1 in a Ras1-independent fashion. In other words, low pH inhibits cAMP synthesis by Cyr1 independently of Ras1 to negatively regulate hyphal growth [53]. However, how the catalytic activity of Cyr1 is downregulated by low pH to reduce cAMP synthesis remains unknown. The identification of Rim20 as a novel Cyr1-interacting protein in this study offers a possible molecular mechanism to solve this mystery. Rim20 participates in the pH response pathway and binds to the transcription factor Rim101 to facilitate the proteolytic activation of Rim101 in alkaline pH [41]. Rim20 may bind to Cyr1 in low pH and inactivate its catalytic activity by proteolytic cleavage similarly.

### 4.6. Mitochondrial proteins as Cyr1 interactors

Four mitochondrion-related proteins, including Atp3, Atp16, Sdh12, and Hem15, have also been identified as novel interactors of Cyr1 in this study. Atp3 and Atp16 are subunits of the mitochondrial ATP synthase localized to the mitochondrial inner membrane [54]. Sdh12 is the homologue to yeast Sdh1, a flavoprotein succinate dehydrogenase coupling the oxidation of succinate to the transfer of electrons to ubiquinone as part of the TCA cycle and the mitochondrial respiratory chain [55]. Hem15 is a mitochondrial inner membrane protein participating in heme biosynthesis [56].

Mitochondria are the primary sites for ATP production and function as the powerhouse of eukaryotic cells. In addition to the roles in metabolism and energy production, the mitochondria of *C. albicans* also play a vital role in many aspects of cellular physiology, including hyphal morphogenesis [57,58]. For example, inhibiting mitochondrial function markedly decreased the cellular level of GTP-Ras1 and blocked hyphal growth. Interestingly, Cyr1 has been shown to participate in the control of Ras1-GTP binding in response to decreased mitochondrial activity, although the underlying mechanism remains elusive [59]. The identification of several mitochondrial proteins as Cyr1 interactors points out one possibility that these physical interactions may mediate the coordination between mitochondrial activity and Ras1-GTP binding.

In summary, by using SILAC-based quantitative proteomics, we have identified 36 novel Cyr1-associated proteins. Six actin regulatory proteins may function as either the upstream regulators of Cyr1 to control its cyclase catalytic activity or downstream effectors to organize the actin cytoskeleton for polarized growth. Four cell wall proteins may activate the Cyr1-dependent cAMP pathway to initiate the switch from yeast to hyphal growth in response to external signals such as attacks by the host immune system. The identification of Rim20 may explain how the catalytic activity of Cyr1 is downregulated by low pH. Interaction of four mitochondrial proteins with Cyr1 may coordinate mitochondrial activity with Ras1-GTP binding. Many other proteins identified in this study remain uncharacterized, and the interaction with Cyr1 may provide a starting point towards their functional characterization. Future studies are required to validate the interactions of these potential binding partners with Cyr1 by other independent methods such as co-IP, yeast two-hybrid assays, and in vitro binding assays. It is also important to identify the domains of Cyr1 for each of the interactions. Undoubtedly, establishing a comprehensive interacting network for Cyr1 will greatly advance our understanding of the regulation and function of this import fungal adenylyl cyclase.

## Author contributions

GZ, experimental design, conducting experiments, data analysis, and preparation of the manuscript; SPN, SILAC-based experiments; LMP, biofilm formation assays; JG, GO term analysis; JG, data analysis of SILAC experiments; YW, experimental design, supervision of the project, writing of the manuscript, and securing funding.

## Acknowledgments

We thank Yan-Ming Wang and Fong Yee Chan for the general technical assistance. We also thank members of the Wang lab for critical reading of the manuscript. This work was funded by the Agency for Sciences, Technology and Research of Singapore with OFIRG/0055/2019 awarded to YW and BMRC042 awarded to GZ.

---

**Figure S1:**
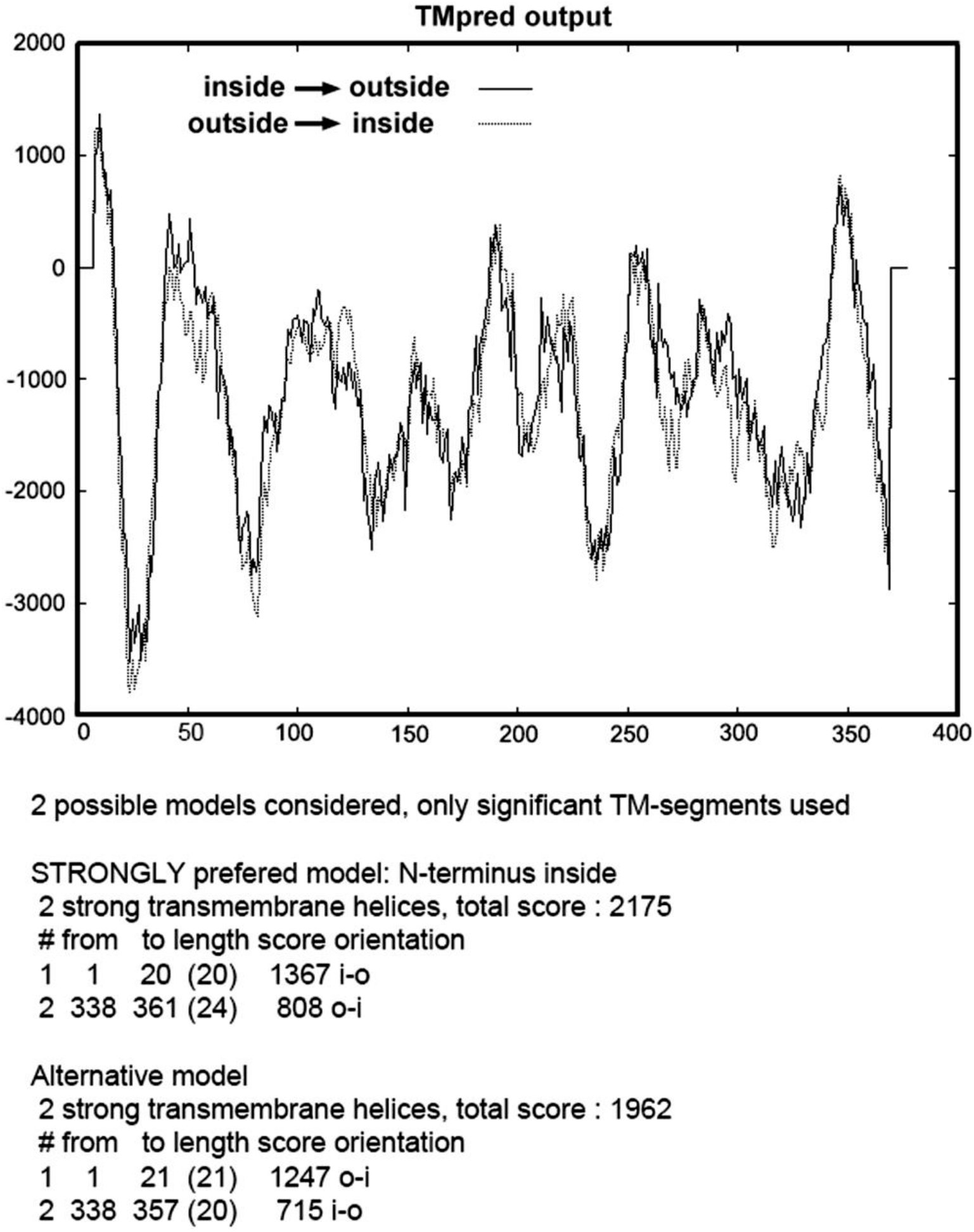
Highly possible transmembrane domains in Mp65 identified by the transmembrane topology analysis. Regions exposed to both the outside and inside of the plasma membrane are listed.

